# Metal Uptake Systems Underpin *Enterococcus faecalis* Virulence in Both Non-Diabetic and Diabetic Wound Infection Models

**DOI:** 10.1101/2025.10.03.680211

**Authors:** Debra N. Brunson, Ling Ning Lam, Shivani Kundra, Shannon M. Wallet, José A. Lemos

## Abstract

Wound infections remain an important medical problem, which is aggravated by the prevalence of multidrug resistant bacteria. Among them, *Enterococcus faecalis* is a major pathogen of surgical site incisional and of diabetic chronic wounds, but factors driving its colonization and persistence in wounds remain poorly understood. Iron, manganese, and zinc are essential cofactors in cellular processes, prompting the host to restrict their availability through mobilization of metal-sequestering proteins, a defense known as nutritional immunity. Previously, we showed that *E. faecalis* strains lacking key iron (Δ5Fe), manganese (Δ3Mn), or zinc (Δ2Zn) uptake systems have impaired virulence. Here, we used an excisional wound model in normoglycemic (C57Bl/6J or B6) and diabetic (C57Bl/6J lepR⁻/⁻ or DB) mice to examine the role of these metal import systems in wounds. The strong upregulation of metal import genes and reduced wound colonization by Δ3Mn, Δ5Fe, and Δ2Zn strains in B6 mice indicate that iron, manganese, and zinc are limited during wound infection. While Δ2Zn and Δ3Mn strains showed no improved colonization in diabetic wounds, the Δ5Fe exhibited a temporary colonization advantage over non-diabetic mice. Quantifications of metal-sequestering proteins lactoferrin, transferrin, calprotectin and psoriasin from intact skin and infected wounds indicated that nutritional immunity, especially iron restriction, is delayed in diabetes. In conclusion, this study underscores the crucial role of trace metal acquisition in *E. faecalis* wound colonization and suggests differences in metal bioavailability between diabetic and non-diabetic wounds, helping to explain the increased susceptibility of diabetic wounds to chronic infection.

## INTRODUCTION

Wound infections cover a range of soft tissue complications, including those resulting from incisional surgery, burns, and traumatic injuries. Estimates for yearly global wound care expenditure is almost $300 billion and with an expansion in the aging populations and co-morbidities such as diabetes and hypertension, this number is only expected to increase over time (1, 2). In healthy individuals, the wound healing process follows four tightly-controlled overlapping stages: (i) hemostasis; which occurs immediately after injury and leads to the formation of blood clots, (ii) inflammation; which occurs 1-3 days after injury and involves clearance of any bacteria present and necrotic debris to prepare the wound bed for new growth, (iii) proliferation; which can last days or weeks and includes re-epithelialization, restoration of the vasculature, and formation of granulation tissue, and (iv) remodeling; which can take several weeks and even years and involves the development of tensile strength of the new tissue and complete closure of the wound (3, 4). This process depicts what typically occurs in a normal healing wound; however, due to internal and/or external factors, some wounds stall in their progression and become chronic non-healing wounds, commonly associated with unresolved inflammation, elevated levels of reactive oxygen species, limited blood circulation, and often times, become infected with opportunistic wound pathogens such as *Staphylococcus aureus*, *Pseudomonas aeruginosa, Escherichia coli* and *Enterococcus faecalis* (2, 5). One of the most common and difficult types of chronic wounds to treat are “diabetic wounds”, notably diabetic foot ulcers, typically associated with Type II Diabetes (T2D) or insulin resistance (6–8). These infections can often lead to lengthy hospital stays, amputations, sepsis, and death (7). As such, a better understanding of both non-diabetic and diabetic wound environments and the underlying mechanisms that dictate host-pathogen interactions can pave the way for the development of new therapies to prevent and treat wound infections.

Of interest, an important host defense mechanism used to combat infections is nutritional immunity, which stems primarily from innate immune cells secreting copious amounts of proteinaceous metal chelators that starve invading pathogens of trace metals such as iron, manganese, and zinc (9–11). These proteins include iron binding proteins ferritin, which sequesters intracellular iron; transferrin, which binds iron in circulation (9, 12), lactoferrin, which restricts access to iron at mucosal surfaces and modulates the host inflammatory response (13–16), and lipocalin-2, which targets iron complexed to bacterial siderophores (17, 18).

Additionally, S100A8/A9 or calprotectin, primarily produced and released by neutrophils at the site of infection, avidly binds divalent metals and is particularly known for depriving pathogens of manganese and zinc, while S100A7 (psoriasin), abundant in the epidermis, contributes to host defense through zinc sequestration (19–21). To overcome metal starvation, bacterial pathogens evolved sophisticated methods to maintain metal homeostasis that include expression of surface-associated high-affinity metal transporters, secretion of metallophores, and, eventually, activation of alternative metal-sparing enzymes (9, 22).

While our understanding of nutritional immunity and trace metal homeostasis in healthy (normoglycemic) individuals is relatively well understood (9, 16, 23–27), fundamental knowledge of these processes in the context of uncontrolled diabetes is limited. In agreement with clinical observations, prior studies that have used diabetic wound infection mouse models have shown that prevalent wound pathogens including *S. aureus*, *P. aeruginosa*, and *Streptococcus agalactiae* maintain significantly higher bacterial burdens in wounds of diabetic mice when compared to normoglycemic mice (28–31). These studies support the notion that higher bacterial titers are associated with an immunocompromised, nutrient-rich diabetic host environment that is less capable of restricting microbial growth (28, 30, 32–34). Along these lines, it is tempting to speculate that uncontrolled diabetes could also impair nutritional immunity activation in diabetic wounds facilitating bacterial proliferation. In fact, a recent study has showed that *S. agalactiae* genes encoding zinc, manganese, and nickel transporters were highly expressed in infected wounds of non-diabetic mice compared to diabetic mice wounds (32). In a subsequent study, this same group demonstrated that the loss of these transporters impaired *S. agalactiae* colonization in non-diabetic wounds but had no effect in diabetic wounds, despite the higher levels of calprotectin present in diabetic wounds (28).

The World Health Organization has listed *E. faecalis*, a major nosocomial pathogen, as the 4^th^ most common pathogen in surgical site infections (35). In addition, *E. faecalis* is amongst the most frequently isolated bacteria from diabetic wounds (36–39), and has been associated with a high degree of antibiotic treatment failure and poor outcomes that includes progression to bone infection, eventual need for limb amputation, and sepsis-mediated death (37, 40, 41).

Despite its high prevalence in wound infections, our understanding of the factors that promote *E. faecalis* colonization and persistence within the wound environment is still limited. A mouse model of enterococcal wound infection has demonstrated that *E. faecalis* modulates the wound micro-environment to evade immune clearance and delay healing (42). Furthermore, the ability of *E. faecalis* to undergo *de novo* purine biosynthesis and metabolize mannose and galactose has been shown to contribute to its capacity to colonize and persist within wounds (43). While previous works have focused on the microbial factors and host responses that contribute to wound infections, the importance of nutritional immunity and its possible association with chronic diabetic *E. faecalis* wound infections have not been investigated. For the past several years, our group has shown that the ability to acquire trace metals, iron, manganese, and zinc is critical for the virulence of *E. faecalis* (44–46). Here, we leveraged the availability of well-characterized *E. faecalis* mutants with major defects in manganese (46), iron (44), or zinc uptake (45) to assess their ability to colonize wounds in both normoglycemic and diabetic mice. While our findings underscore the importance of trace metal acquisition in *E. faecalis* wound infections and reveal nuanced differences in the expression of host-derived proteinaceous metal chelators in a T2D mouse model, a more definitive correlation between impaired nutritional immunity and chronic colonization of diabetic wounds by *E. faecalis* remains to be established.

## RESULTS

### Iron, manganese, and zinc uptake systems of *E. faecalis* are critical for wound colonization

We previously showed that the *E. faecalis* core genome encodes four iron ABC-type transporters, FeoAB, FhuDCBG, FitABCD and EmtABC, as well as a dual iron/manganese ABC-type transporter, EfaCBA (44, 47). We also identified and characterized two manganese NRAMP-type transporters, MntH1 and MntH2, and an ABC-type zinc transporter that utilizes two interchangeable substrate-binding proteins, AdcA and AdcAII, which function with the shared AdcB permease and AdcC ATPase subunit (44, 45, 47). Using the invertebrate model *Galleria mellonella* and mammalian infection models (mouse and rabbit), we demonstrated that strains of *E. faecalis* with major defects in manganese (Δ3Mn), zinc (Δ2Zn), or iron (Δ5Fe) uptake exhibit significantly reduced virulence across all tested hosts (44, 45, 47). While previous work using the excisional wound mouse model showed that the Δ5Fe strain colonized wounds poorly (44), the roles of manganese or zinc acquisition during *E. faecalis* wound infection were not evaluated in those studies.

Here, we assessed the ability of the Δ3Mn and Δ2Zn strains to colonize excisional wounds in C57BL/6J (B6) mice and included the Δ5Fe strain to directly compare the relative importance of each metal during infection. In addition to examining the acute phase of infection (1- and 3-days post-infection), we also quantified bacterial colonization 7 days post-infection to evaluate the potential for chronic wound colonization. While there were no differences in the bacterial burden of wounds infected with the parent strain (OG1RF) and any of the individual metal uptake mutant at 1-day post-infection, all mutants were recovered at significantly lower levels 3 days post-infection. Among them, the Δ2Zn strain exhibited the smallest reduction (∼1-log) compared to OG1RF, whereas the Δ3Mn strain showed the greatest decrease (∼1.8-log) in bacterial load (**Fig. 1**). By day 7 post-infection, the difference in bacterial titers between wounds infected with the Δ2Zn strain and those infected with the parent strain was no longer significant while both Δ3Mn and Δ5Fe continued to be recovered at significantly lower levels (∼1-log) compared to OG1RF (**Fig. 1**).

**Figure 1.**
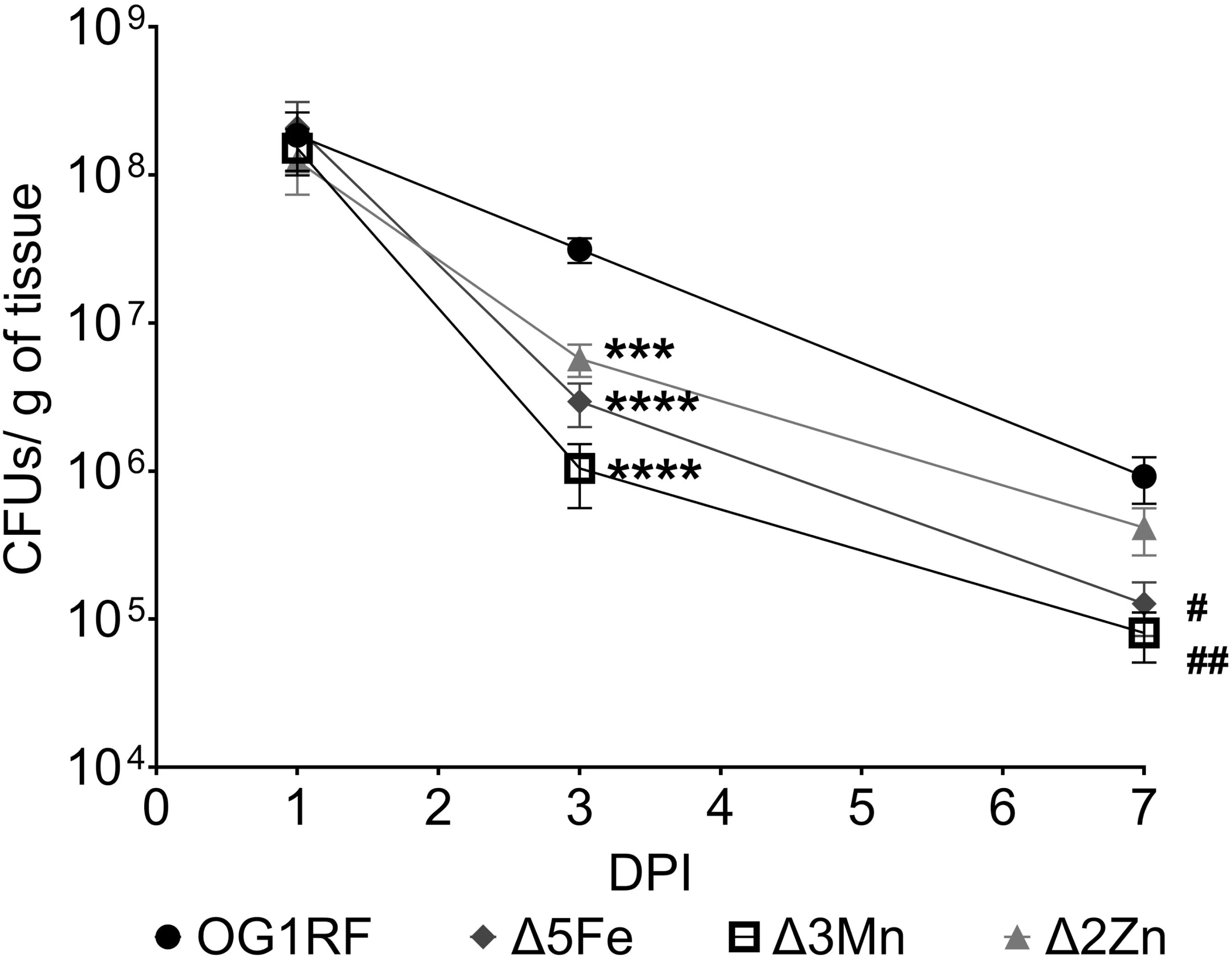
Seven-week-old B6 mice were wounded with a 6 mm biopsy punch and infected with 10^8^ CFU of either *E. faecalis* OG1RF, Δ5Fe, Δ3Mn, or Δ2Zn strains. Wounds were excised at 1-, 3-, and 7-days post-infection (p.i.) and homogenized in PBS for CFU determination by plating on BHI agar. Each point represents the average and standard deviation from at least 9 infected mice. Outliers were removed with a ROUT standard outlier test. Statistical significance compared to the parental strain was determined using a Mann-Whitney test. Day 3: \*\*\**p*≤0.001, \*\*\*\**p*≤0.0001; Day 7= #p≤0.05, ##p≤0.01.

Collectively, the results described above are aligned with previous studies proposing that essential trace metals are not readily bioavailable in wound microenvironments (28, 32, 43). To support this observation, we compared the transcriptional levels of *E. faecalis* genes involved in the uptake of iron (*fitD*, *emtC*, *feoB*, and *fhuG*), manganese (*mntH1* and *mntH2*), both manganese and iron (*efaA*), and zinc (*adcA*, *adcAII*) between cells grown in brain heart infusion (BHI) medium and those recovered from infected wounds of B6 mice on days 1, 3 or 7. Notably, we have consistently shown that BHI contains low concentrations of iron (∼5 µM), manganese (<1 µM), and zinc (∼10 µM) (44, 46, 48), and thus cannot be considered a metal-rich medium.

Consistent with our expectations, transcription of iron, manganese and zinc uptake genes was significantly elevated in bacteria from infected wounds relative to BHI-grown counterparts (**Fig. 2A-C**). We also assessed the expression of *dpr* and *sodA* as markers of iron overload (*dpr*) and oxidative stress (both genes) in infected wound samples. Consistent with the upregulation of iron transporters, an indicative of iron starvation, *dpr*, which encodes the major iron storage protein, was significantly downregulated (**Fig. 2D**). In contrast, *sodA* transcription was modestly upregulated in the wounds, suggesting that *E. faecalis* may also experience ROS stress in the wound microenvironment. (**Fig. 2D**).

**Figure 2.**
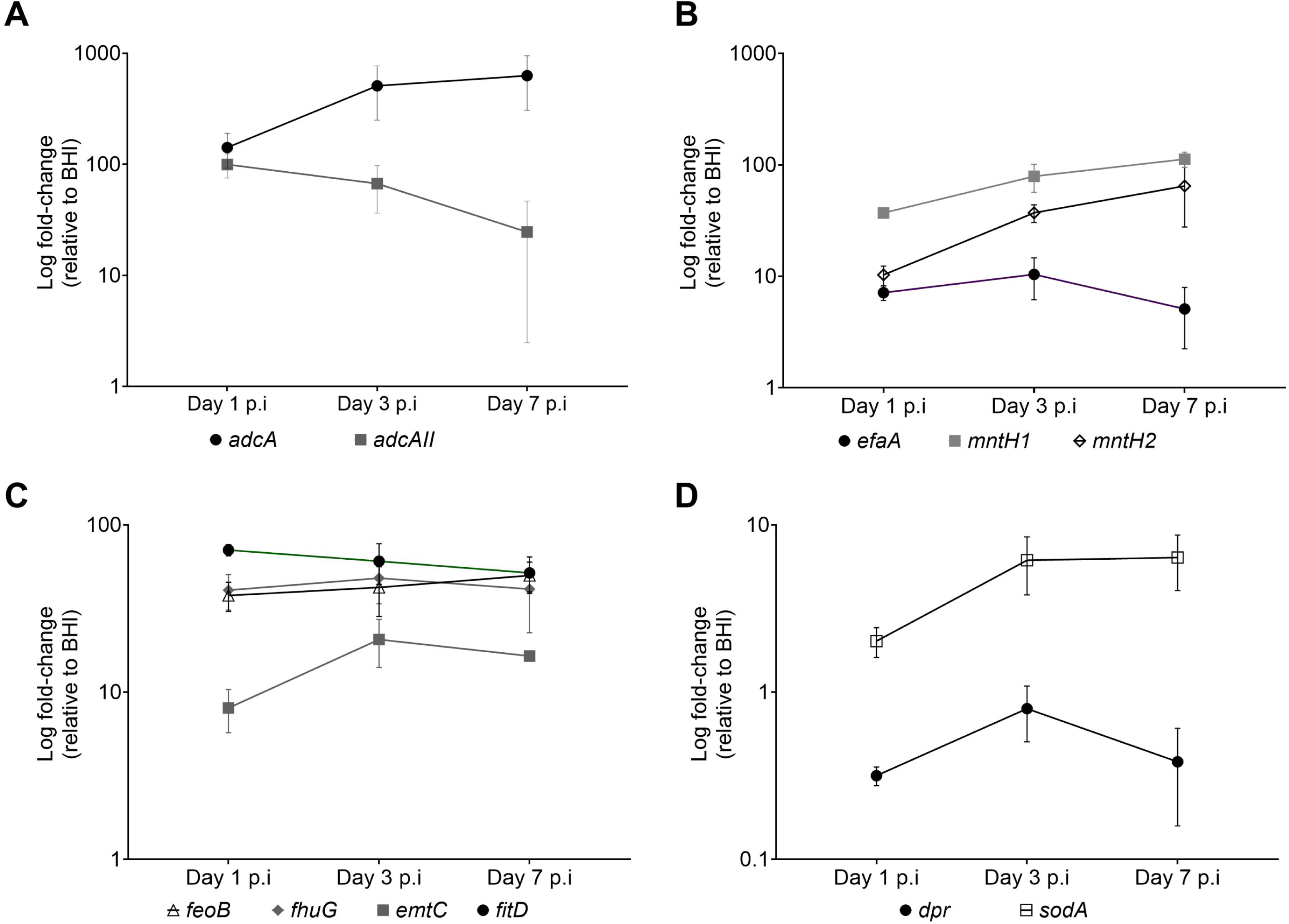
Transcriptional profile (relative to BHI) of *E. faecalis* metal transporters in infected wounds of B6 mice at 1-, 3-, and 7-days post-infection. (A) zinc transporters (*adcA* and *adcAII*), (B) manganese transporters (*efaA*, *mntH1*, and *mntH2*), (C) iron transporters (*feoB*, *fhuG*, *emtC*, and *fitD*), and (D) iron storage (*dpr*) and oxidative stress management (*sodA*). Statistical significance was determined using ordinary one-way ANOVA Fisher’s LSD test. \**p*≤0.05, \*\**p*≤0.01, \*\*\**p*≤0.001, \*\*\*\**p*≤0.0001.

### *Enterococcus faecalis* maintains higher bacterial burden in diabetic wounds than in non-diabetic wounds

Previous studies have demonstrated that diabetic mice exhibit higher bacterial burdens in infected wounds compared to normoglycemic mice (28–30). These authors proposed that, in addition to impaired immune responses, the nutrient-rich environment of diabetic wounds significantly contributes to this phenomenon (28, 30, 32–34). Here, we investigated whether *E. faecalis*, like other prevalent wound pathogens, exhibits enhanced infection in diabetic wounds compared to non-diabetic wounds. To address this, we compared *E. faecalis* OG1RF bacterial burden at 1-, 3-, and 7-days post-infection in wounds of B6 (normoglycemic) mice and C57Bl/6J *lepR*^⁻/⁻^ (DB) mice, which recapitulates the physiological and pathological features of T2D in humans (49). Compared to B6 mice, DB mice exhibited significantly higher bacterial titers (∼1-log;) in wounds at 1- and 3-days post-infection (**Fig. 3A**). This difference became much greater by day 7 (∼3-log), as B6 mice continued to clear the infection between days 3 and 7, whereas the bacterial burden in DB mice remained unchanged. Moreover, the infected wounds of DB mice not only failed to heal but increased in size by ∼30% between days 3 and 7, whereas wounds of B6 mice showed ∼50% reduction in size over the same period (**Fig. 3B**). Visual examination revealed progressive healing of infected wounds in B6 mice over time, whereas wounds in DB mice appeared highly inflamed and exhibited extensive accumulation of cellular exudates (**Fig. 3C**).

**Figure 3.**
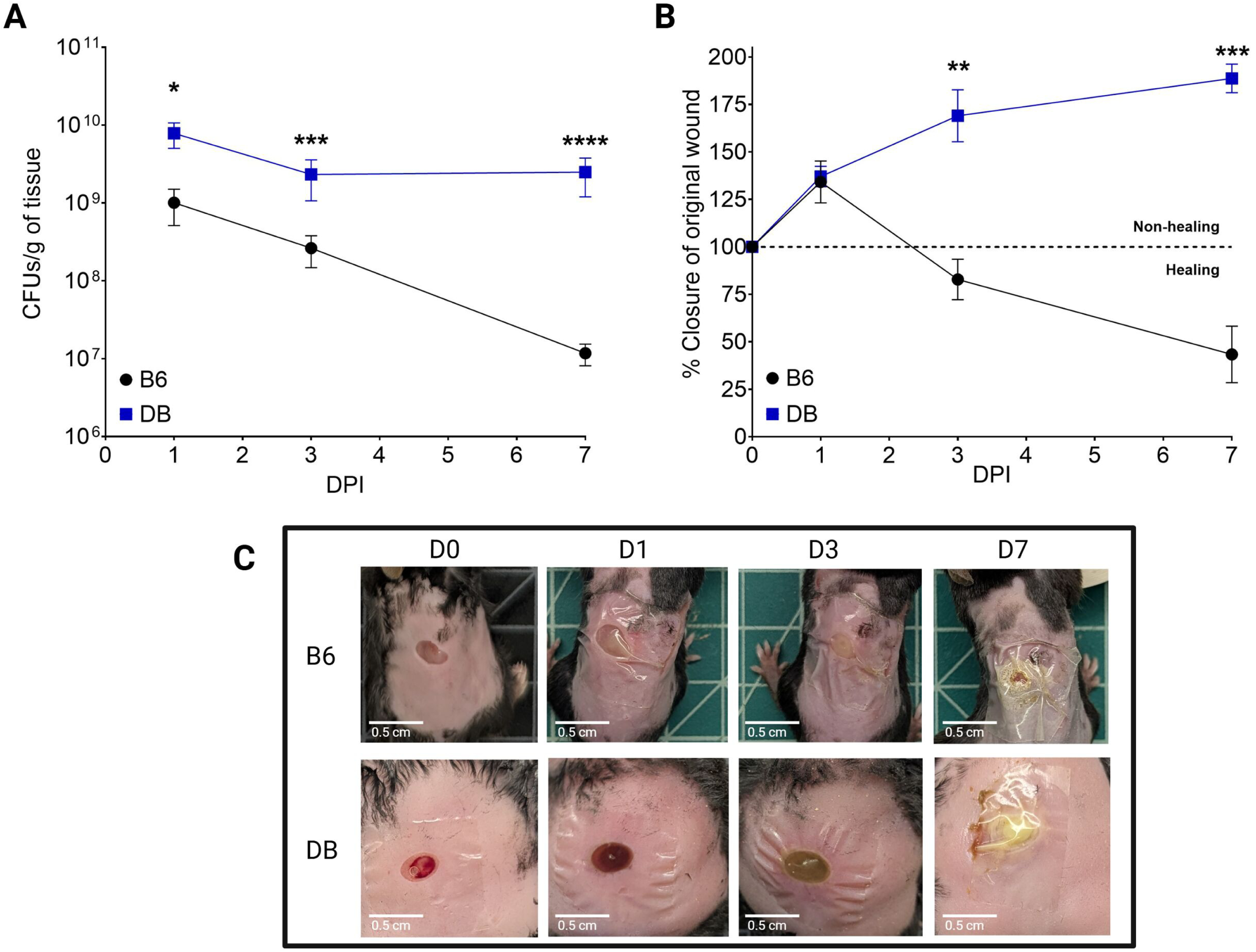
Seven-week-old B6 or DB mice were wounded with a 6 mm biopsy punch and infected with 10^8^ CFU of *E. faecalis* OG1RF. (A) Wounds were excised at 1-, 3-, and 7-days post-infection, homogenized and bacterial CFU determined by plating on BHI agar. (B) Wound area was measured using ImageJ and percent closure determined by comparing to initial wound area. (C) Images of wounds immediately after wounding (D0) and at 1-, 3-, and 7-days post-infection. Each time point represents the average and standard deviation from at least 5 infected mice. Outliers were removed using a ROUT outlier test. Statistical significance was determined using a Mann-Whitney test at each time point. \**p*≤0.05, \*\**p*≤0.01, \*\*\**p*≤0.001, \*\*\*\**p*≤0.0001.

### Trace metal acquisition facilitates *E. faecalis* colonization of both normoglycemic and diabetic wounds

Similar to our findings (**Fig. 2**), a group at the University of Colorado previously demonstrated that *S. agalactiae* strains lacking either manganese or zinc uptake systems exhibit impaired wound colonization in normoglycemic mice (28). They also demonstrated that the impaired wound colonization of metal transport mutants was no longer observed in streptozotocin (STZ)-induced hyperglycemic mice, a model commonly used to study type 1 diabetes (T1D) (28). To further probe the role of iron, manganese, and zinc acquisition in *E. faecalis* colonization of diabetic wounds in DB (T2D) mice, we performed a competitive wound colonization experiment to directly compare relative virulence potential of parent and mutant strains in both B6 and DB mice. Briefly, wounds created in either B6 or DB mice were infected with an inoculum containing equal proportions of each strain (OG1RF, Δ5Fe, Δ3Mn, and Δ2Zn) and relative abundance of each strain recovered from wounds determined at 3- and 7-days post-infection. Regardless of the mouse background, all three mutants were outcompeted by the parental strain at both 3- and 7-days post-infection (**Fig. 4A-B**). Consistent with the single-strain infection experiments in B6 mice (**Fig. 2**), loss of the zinc uptake system (Δ2Zn) had the least impact on colonization fitness whereas disruption of manganese uptake (Δ3Mn) had the most pronounced effect. When compared to B6-infected mice, the Δ5Fe strain was recovered at slightly higher rates (*p*=0.0449) in wounds of the DB mice on day 3; however, this apparent fitness advantage did not persist by day 7. Collectively, these findings validate the critical role of trace metal acquisition in *E. faecalis* wound colonization but do not support that impaired nutritional immunity alone explains the increased permissiveness of the diabetic wound environment to infection.

**Figure 4.**
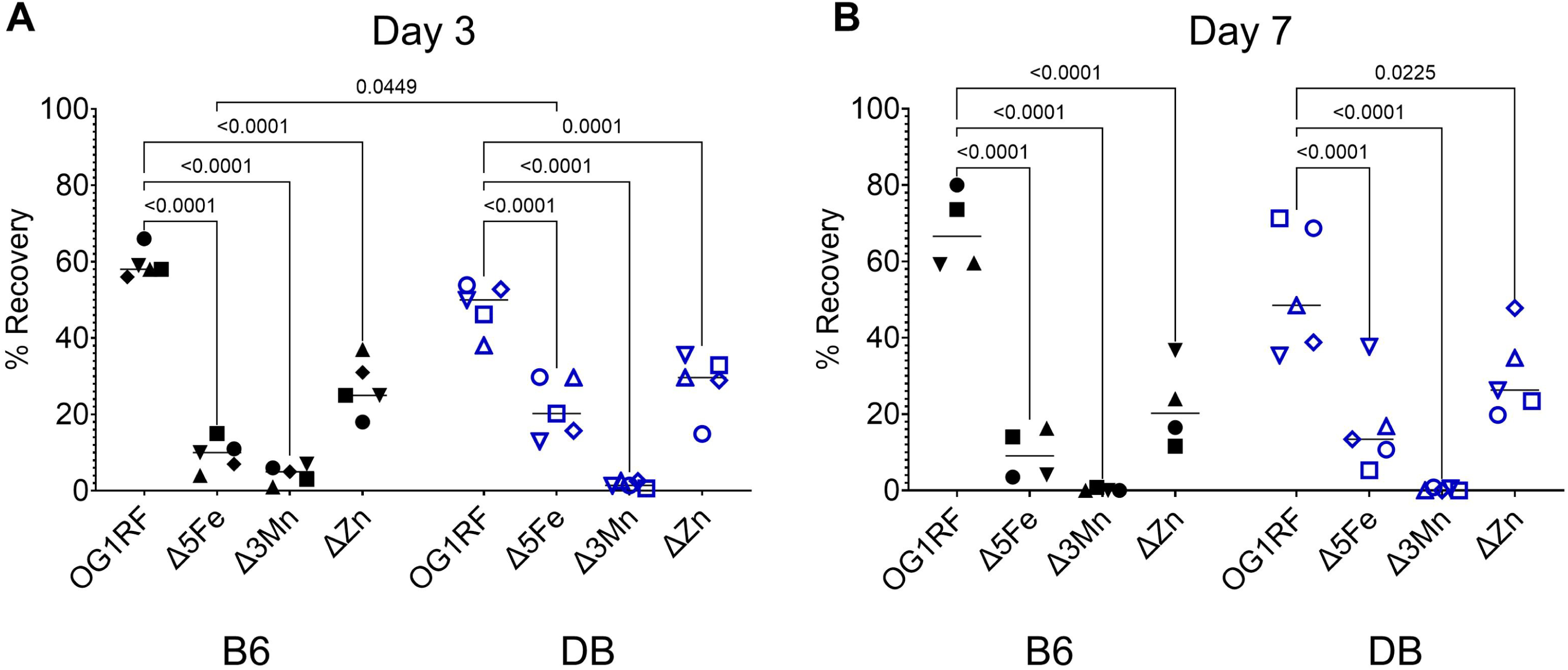
Competitive wound colonization. B6 and DB mice were infected with an inoculum of 10^8^ CFU containing an equal ratio of the *E. faecalis* OG1RF, Δ5Fe, 3Mn, and Δ2Zn strains. At 3- and 7-days post-infection, wounds were excised and homogenized in PBS for CFU determination by plating on BHI agar. Bacterial colonies were screened using gene-specific primers (Table 2). At least 100 colonies were screened from each animal to calculated the percentage recovery of each strain. Statistical significance was determined using a two-way ANOVA with a Šidák’s multiple comparison test. *p* values < 0.05 were considered statistically significant and the exact values are shown in the figure.

### Metal homeostasis may differ between diabetic and non-diabetic wounds

The observation that the Δ5Fe strain was recovered at a modest, yet significantly higher rate in DB wounds on day 3 (**Fig. 4A**) led us to suspect that trace metal bioavailability in the diabetic wound microenvironment may be temporally altered compared to non-diabetic wounds, with greater access to iron in the early stages of healing that diminishes over time. One possibility is that the production and mobilization of host-derived metal chelators such as proteins from the S100 family, transferrin and lactoferrin (9, 20) to the site of infection differ in diabetic wounds when compared to non-diabetic wounds. To test this, we quantified total transferrin (tTF), apo-transferrin (aTF), lactoferrin, calprotectin (CP/S100A8/A9), and psoriasin (S100A7) levels in skin tissues harvested from naïve and OG1RF-infected wounds in both B6 and DB mice.

Interestingly, transferrin levels were approximately 10-fold higher in naïve skin tissue of B6 mice compared to DB mice; however, this difference was no longer evident at any time point in the infected wounds (**Fig. 5A**). Similarly, apo-transferrin levels were approximately two-fold higher in naïve skin tissue of B6 mice compared to DB mice but were markedly reduced upon infection in both groups (**Fig. 5B**). This supports the notion that transferrin plays an active role in the host’s nutritional immunity strategy. Nevertheless, comparison of the aTF-to-tTF ratio in B6 and DB mice revealed a slightly different pattern. Specifically, this ratio was approximately 3.5-fold higher in the naïve skin tissue of DB mice compared to B6 mice but declined by about 50% after one day of infection. By days 3 and 7, no statistically significant differences were observed between the groups (**Fig. 5C**).

**Figure 5.**
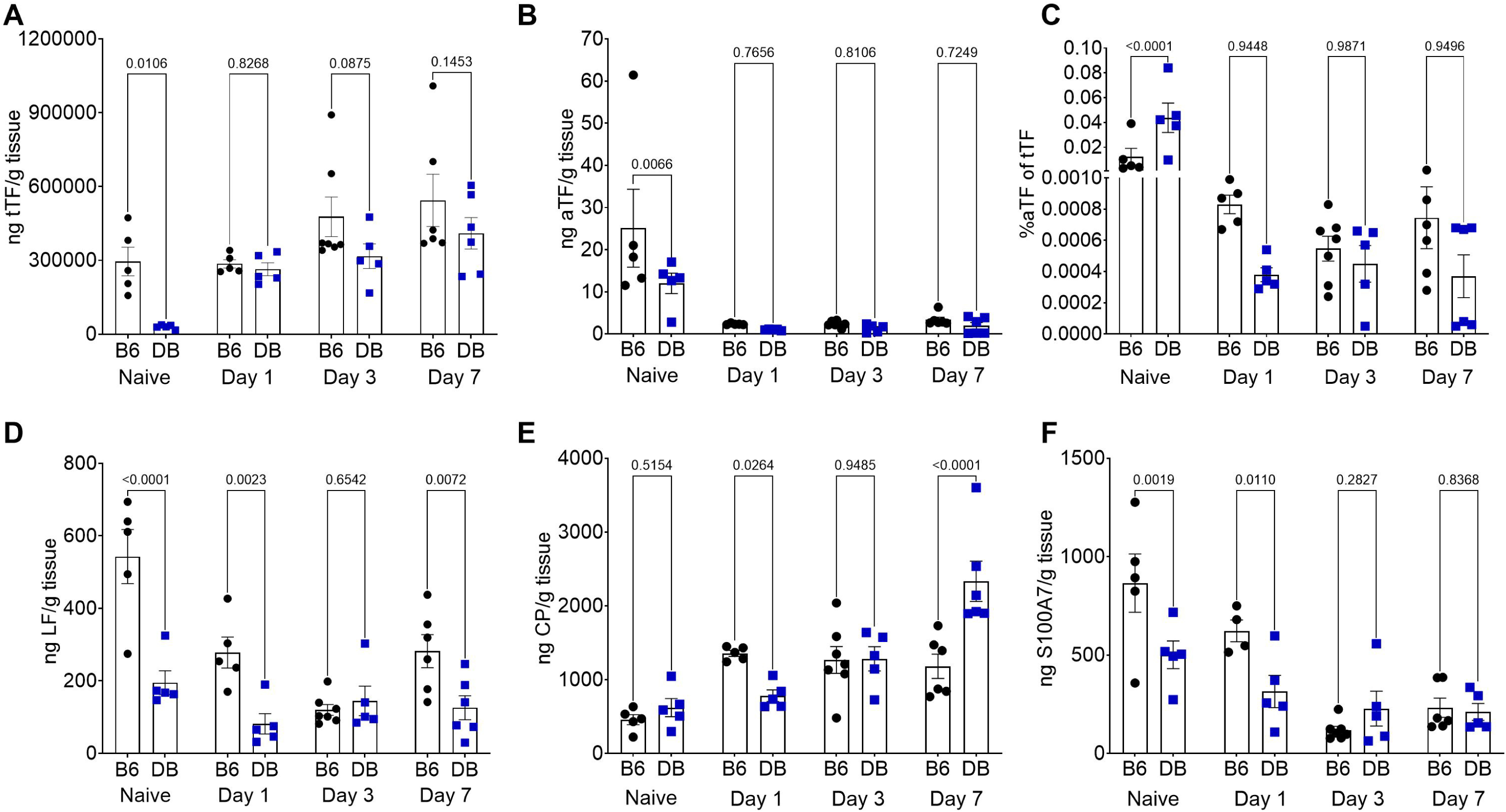
Abundance of host metal chelators in naïve skin and infected wounds of B6 and DB mice. Naïve skin from biopsy punch and excised wounds from 1-, 3-, and 7-days post-infection were homogenized in metal-free PBS containing protease inhibitor. ELISA kits were used to determine the abundance of each host metal chelator and normalized to grams of tissue. (A) Total transferrin (tTF), (B) apo-transferrin (aTF), C) percentage aTF over tTF (%aTF=(aTF/tTF) x 100%, (D) lactoferrin (LF), (E) calprotectin (CP), and (F) S100A7 (psoriasin). Each data point represents the value from a single animal with at least 5 animals used. Statistical significance was determined using a two-way ANOVA with a Šidák’s multiple comparison. *p* values < 0.05 were considered statistically significant and the exact values are shown in the figure.

**Figure 6.**
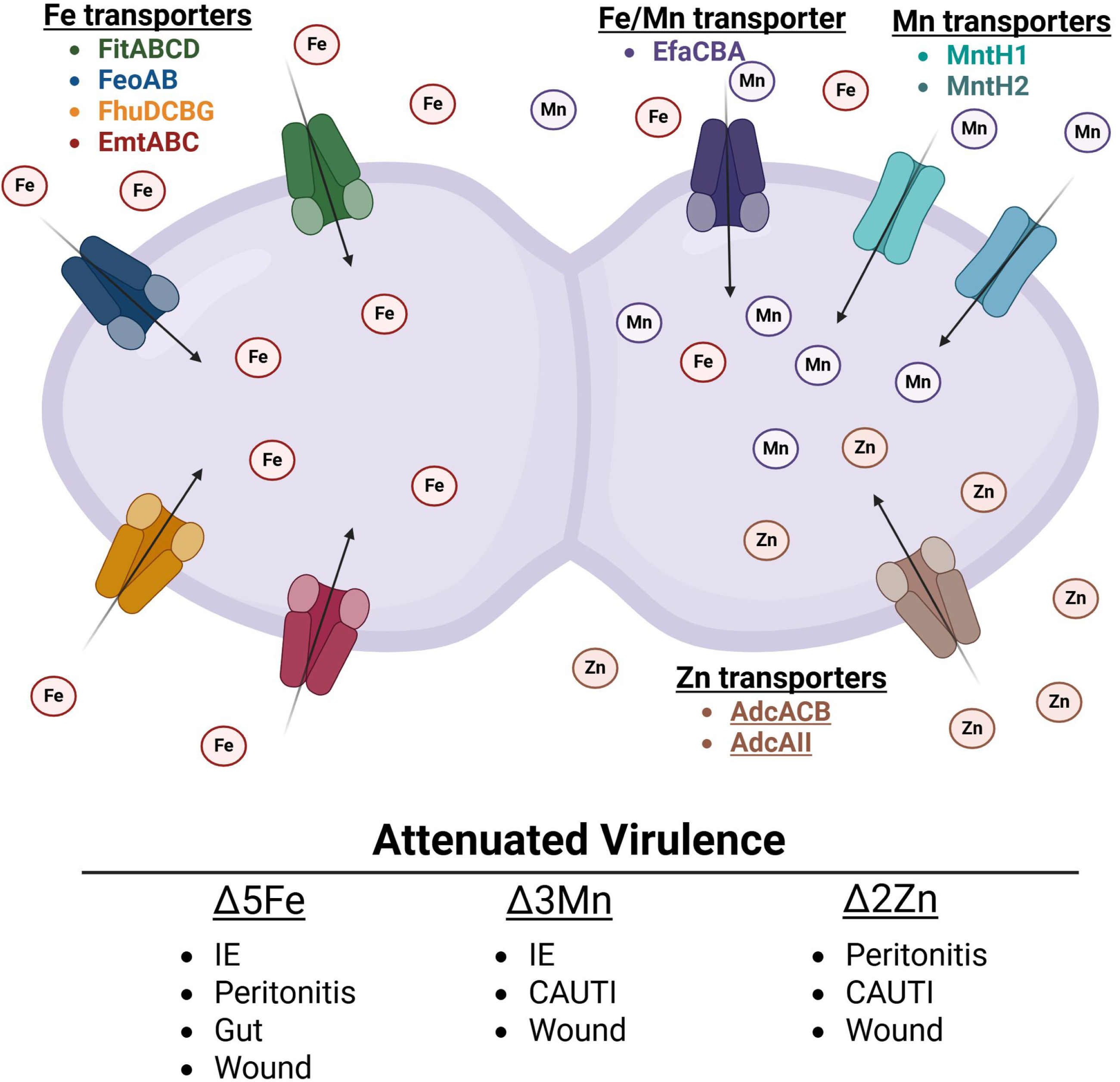
High-affinity iron, manganese and zinc transporters promote *E. faecalis* virulence in systemic and localized infections.

Lactoferrin levels followed a similar pattern to total transferrin, with levels approximately 2.5-fold higher in B6 naïve skin tissue compared to DB mice. This difference persisted at day 1 (∼3-fold) and day 7 (∼2-fold) post-infection but was not observed on day 3 (**Fig. 5D**). While calprotectin levels were similar in naïve skin tissue of B6 and DB mice, infected B6 wounds showed significantly higher calprotectin levels (∼50%) at 1-day post-infection compared to DB mice. Notably, calprotectin levels remained relatively stable over the 7-day infection period in B6 mice, whereas they increased steadily in DB mice, rising approximately 2.5-fold between days 1 and 7 post-infection (**Fig. 5E**). By day 7, calprotectin levels were nearly twice as high in DB mice compared to B6 mice. Considering that calprotectin is produced in large quantities by neutrophils and other immune cells in response to infection and is used as a gold standard marker for inflammation (20), this latter result was expected, given the higher bacterial burden and the highly inflammatory appearance of DB-infected wounds at day 7. Finally, psoriasin levels were approximately twofold higher in naïve skin tissue and at one day post-infection in B6 mice compared to DB mice (**Fig. 5F**). However, by days 3 and 7 post-infection, psoriasin levels declined to comparable levels in both mouse strains.

## DISCUSSION

It is estimated that one third of the bacterial proteome requires a metal cofactor (50), making the ability of bacteria to maintain an adequate intracellular supply of biometals essential for bacterial fitness (10). Accordingly, metal transporters (both import and export systems), metal chaperones, and metal storage proteins are consistently linked to bacterial virulence (10, 22, 51–53). Using an excisional wound mouse model, we confirmed previous findings that the loss of multiple iron transporters impairs wound colonization and extended this observation to include manganese and zinc transporters. In addition, we demonstrate that diabetes greatly increases host susceptibility to enterococcal wound infection during both the acute and chronic phases.

While the roles of iron, manganese, and zinc import in *E. faecalis* virulence are relatively well defined (44–46), the factors regulating the expression and activity of individual trace metal transporters, particularly within the wound microenvironment, remain poorly understood.

Previous studies have shown that transcription of the N-ramp manganese transporter *mntH2* and dual iron and manganese ABC transporter (*efaCBA*) are strongly induced when *E. faecalis* is grown *ex vivo* in human blood or human urine, as well as *in vivo* using an intra-peritoneal challenge (peritonitis) mouse model (54–56). Furthermore, our lab has shown that the three high-affinity transport systems associated with manganese uptake are differentially regulated in two different animal models (46). Specifically, we showed that *efaA* is highly upregulated, the N-ramp transporter *mntH1* is downregulated, while *mntH2* remains unchanged in the IE rabbit model. On the other hand, *mntH2* is highly upregulated and *efaA* and *mntH1* remain unchanged in the CAUTI mouse model. In contrast, we show here that *mntH1* is the most highly upregulated manganese transporter in infected wound beds, closely followed by *mntH2*, with *efaA* showing only a modest increase. This is also in stark contrast to what is observed *in vitro* where *efaA* and *mntH2* transcription greatly increases after manganese starvation but not *mntH1* (57). While we lack information on how zinc transporters are regulated *in vivo*, we have shown previously that both *adcABC* and *adcAII* are repressed under high zinc conditions and strongly induced after treatment with the zinc-chelating agent TPEN (N,N,N′,N′-tetrakis(2-pyridinylmethyl)-1,2-ethanediamine) (45, 58, 59). Here, we show that transcription of both *adcA* and *adcAII* is strongly induced in wounds one day post-infection. However, while *adcA* transcription declines thereafter, *adcAII* expression continues to rise. These findings suggest that AdcAII becomes the predominant zinc-binding lipoprotein during the chronic stages of wound infection. In addition to the dual iron transporter *efaA*, the other four iron transporters, *fitD*, *emtC*, *feoB*, and *fhuG*, were strongly and consistently upregulated in infected wounds. Together, these results support the idea that these biometals are not readily accessible in the wound microenvironment while also underscoring the importance and complexity of trace metal homeostasis regulation in *E. faecalis*.

To investigate the possible role of nutritional immunity, or a defect thereof, in chronic diabetic wounds, we compared the fitness profiles of iron, manganese, and zinc transport mutants in wounds of both B6 (non-diabetic) and DB (T2D) mice. We also monitored the temporal changes in the levels of major metal-sequestering host proteins, namely transferrin, lactoferrin, calprotectin, and psoriasin, in these wounds. Unlike *S. agalactiae*, in which deletion of either manganese or zinc transporters impaired its ability to colonize wounds of normoglycemic mice but not of diabetic mice (28), *E. faecalis* strains with impaired manganese (Δ3Mn) or zinc (Δ2Zn) uptake exhibited similar infectious potential in either B6 or DB mice. Several, not mutually exclusive, reasons may explain these apparently discrepant results. First, it is conceivable that *E. faecalis* has a higher demand for these metals than *S. agalactiae*. Although less likely given the similar number of high-affinity metal uptake systems shared by the two species (28, 44–46, 60, 61), it is also possible that *S. agalactiae* is simply better equipped to scavenge trace metals than *E. faecalis*. In either case, potential defects in the activation of metal sequestration strategies during diabetes may still be insufficient to support the growth of *E. faecalis* manganese and zinc transport mutants. Additionally, differences in experimental conditions might also have contributed to these differences, particularly considering that we used C57Bl6J lepR^-/-^ mice, which models T2D, whereas the *S. agalactiae* study employed the streptozotocin-induced diabetes model, which more closely resembles T1D (28).

Although the Δ3Mn and Δ2Zn strains do not appear to benefit from the more permissive microenvironment of DB mice wounds, the Δ5Fe strain displayed a slight advantage given its higher recovery rate from DB wounds at 3-days post-infection when compared to B6 mice wounds. In alignment with this observation, both total transferrin and lactoferrin levels were significantly lower in DB naïve skin tissue when compared to B6 mice. In the case of lactoferrin, this difference could be also observed on days 1 and 7 post-infection. Levels of the S100 proteins calprotectin and psoriasin in DB wounds were also significantly lower at 1-day post-infection. Although colonization defects were observed only in the Δ5Fe strain at a single post-infection time point, these findings indicate that normoglycemic mice are better prepared to combat opportunistic wound infections than diabetic mice. This supports the idea that diabetes impairs nutritional immunity, particularly the timely activation of iron-restricting mechanisms.

In conclusion, here we show that the ability to scavenge essential trace metals is critical to *E. faecalis* wound infection. We also show that, as expected, diabetic mice struggle to clear infections and are more susceptible to developing chronic wound infections compared to normoglycemic mice. While this study provides some evidence for impaired activation of nutritional immunity in the context of T2D, based on apparent differences in the surveillance and mobilization of metal-sequestering proteins at infection sites in B6 and DB mice, further research is needed to substantiate this possibility. In particular, studies that directly compare bioavailable metal pools in the skin and wound beds of diabetic versus non-diabetic animals are necessary. Additionally, hyperglycemia has been associated with aberrant protein glycosylation, which may further compromise host immune responses and metal-regulating pathways (62–64). Both transferrin and lactoferrin undergo N-linked glycosylation, and alterations in their glycosylation patterns have been associated with defects in protein stability, trafficking, and activity (65, 66). Prior research has shown that transferrin glycosylation patterns vary among individuals and are influenced by factors such as age, sex, and body weight (65). Building on this, it is plausible to speculate that the hyperglycemic environment characteristic of uncontrolled diabetes could promote aberrant glycosylation of iron-sequestering proteins like transferrin and lactoferrin, potentially impairing their function and thereby compromising the timely activation of nutritional immunity. Beyond this work with *E. faecalis* and previous investigations involving *S. agalactiae* (28), further research is needed to explore the role of trace metal homeostasis in wound infections caused by other prominent pathogens, such as *S. aureus* and *P. aeruginosa*, as well as in polymicrobial infections. In closing, this investigation expands our understanding of host-pathogen interactions in wound infections and identifies differences in metal availability that may inform new therapeutic strategies, particularly for at-risk populations.

## MATERIALS AND METHODS

### Bacterial strains and growth conditions

The strains used in this study are from −80°C freezer stocks previously generated by our lab (44–46). All strains were routinely grown aerobically in brain heart infusion (Difco Laboratories) at 37⁰C. The Δ2Zn strain was supplemented with 100 µM ZnSO_4_ and the Δ3Mn strain was grown with 100 µM MnSO_4_ to overcome growth defects observed in non-supplemented BHI (45, 47). Chemical and biological reagents were purchased from Sigma Aldrich unless stated otherwise.

### Mouse wound excision model

These experiments were performed under protocol 202011154 approved by the University of Florida IACUC. Overnight cultures and starter cultures were obtained as previously (44), with minor modifications. Briefly, the bacterial inoculum was prepared by washing overnight grown cells in 0.5 mM EDTA in PBS once and twice in trace metal grade PBS. For single strain infection, cell pellets were concentrated to 1 x 10^10^ CFU ml^-1^ and stored on ice until use. For competitive index infections, cells were prepared as above, and all four strains were mixed in an equal ratio. Seven-week-old C57BL6J (B6) or B6.BKS(D)-*Lepr*^db^J (DB) mice (transgenic mice of the C57BL6J background with a mutation in the leptin recepto) were purchased from Jackson laboratories and allowed to acclimate at our facilities for at least 3 days. On the day of the experiment, animals were anesthetized using isoflurane, their backs shaved, and the excisional wound created using a 6 mm biopsy punch. Infection of wounds and enumeration of bacterial burden were performed as previously described (42, 44), using tissue homogenate collected at 1-, 3-, or 7-days post infection. For competitive index infections, the inoculum and wound homogenates were plated on selective TSA plates, and PCR was performed to assess the number of each mutant strain recovered using primers specific to iron (*fitD*), manganese (*mntH2*), and zinc (*adcA*) transport (**Table 1**). At least 100 colonies were screened per wound to determine competitive index of each strain.

**Table 1.**
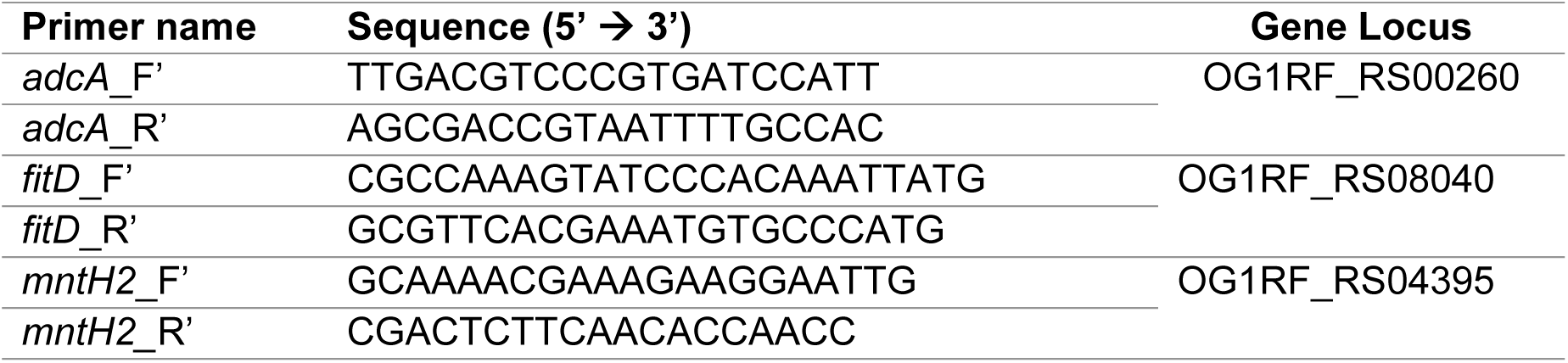
Primers used for PCR-based strain identification in competitive index experiments.

**Table 2.**
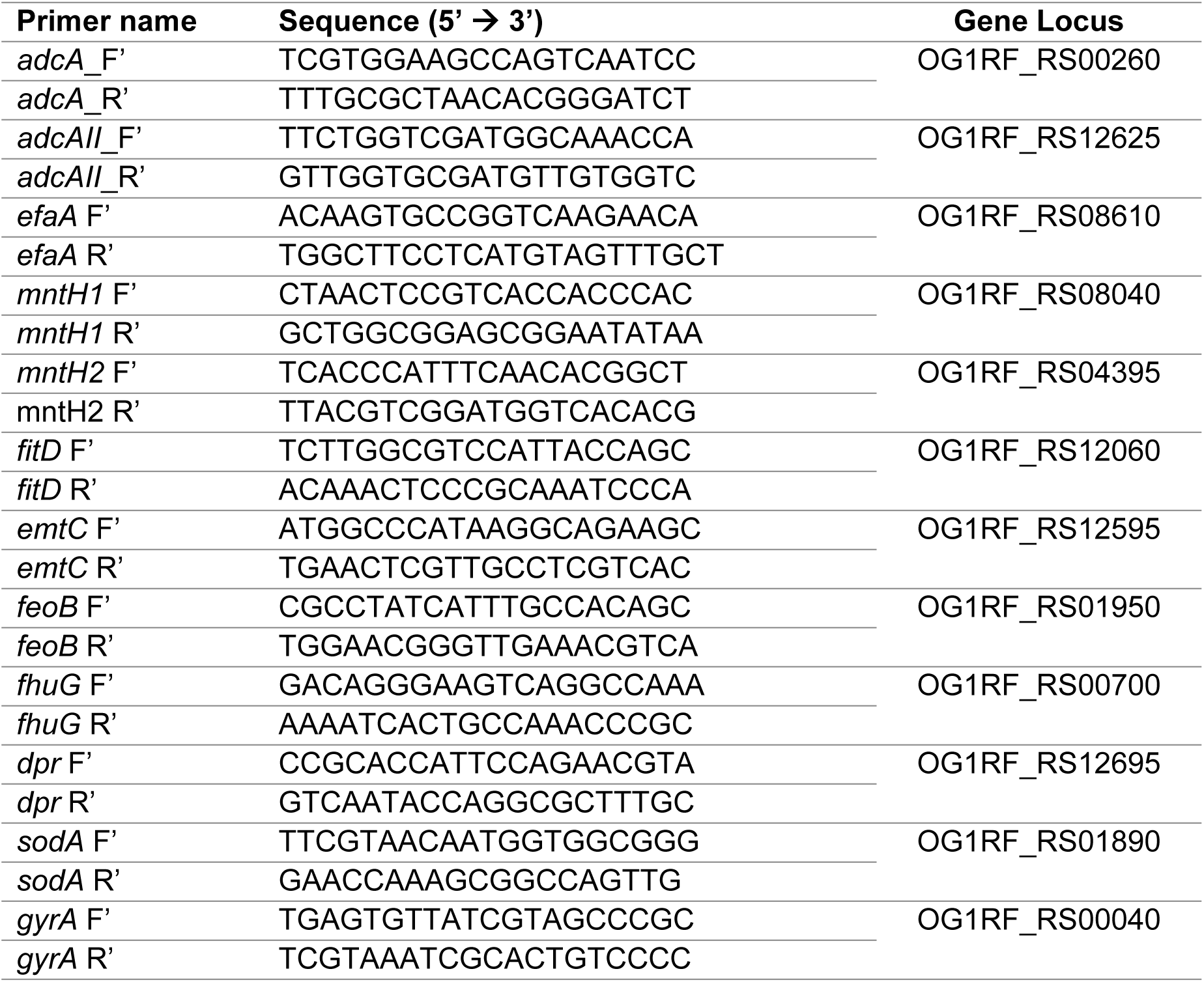
Primers used for qRT-PCR.

### Quantitative real-time PCR

To obtain total RNA from BHI-grown cultures, *E. faecalis* OG1RF was first grown overnight in BHI. Starter culture was normalized to an OD_600_ of 0.5, inoculated at a ratio of 1:20 into fresh BHI and incubated at 37°C for 1 hr. Post incubation, cells were harvested via centrifugation at 4000 rpm for 10 min, washed once with PBS and incubated with lysozyme (Sigma-Aldrich) (20 mg ml^-1^) for 30 min at 37°C. Next, cells were harvested by centrifugation and total RNA extracted using the PureLink RNA minikit (Invitrogen), according to the manufacturer’s instructions. To isolated *E. faecalis* total RNA from infected wounds, skin encompassing the wound site was excised and placed into 2 ml of RNA*later* stabilization solution (Invitrogen) and incubated on ice for 1 hr. The mouse skin was then transferred into 1 mL of TRizol (Life Technologies), processed according to manufacturer’s protocols using the Max Bacterial Enhancement Reagent (Invitrogen), with minor modifications. Briefly, the entire suspension, including the mouse tissue, was homogenized using a hand-held homogenizer (Benchmark Scientific), followed by 3 cycles of bead-beating (40 sec each and chilled on ice for 2 min). Samples were centrifuged for 20 min at 12,000 rpm at 4°C to pellet cell debris, and the aqueous suspension transferred to a new tube. To the homogenized aqueous suspension, chloroform (Sigma-Aldrich) was added, according to TRizol (Life Technologies) Reagent’s standard protocol. The top layer (aqueous phase) containing RNA was transferred to a new tube and total RNA was isolated by ethanol-precipitation. Total RNA was pooled from at least 3 mice to generate one biological replicate (n=3). RNA purification and removal of DNA were performed using a Turbo DNA-free kit (Thermo Fisher). Removal of mammalian RNA and isolation of bacterial RNA were performed using Microb*Enrich* kit (Invitrogen). The tissue extracted bacterial RNA was quality checked using the RNA ScreenTape on a TapeStation instrument (Agilent Technologies) and quantified using Qubit at University of Florida Interdisciplinary Center for Biotechnology Research (ICBR) Gene Expression & Genotyping Core facility. Synthesis of cDNA was performed using a High-capacity cDNA Reverse Transcription Kit (Applied Biosystems). Quantitative real-time PCR (qRT-PCR) was performed using iTaq Universal SYBR supermix (BioRad) with the primers listed in **Table 2**. For quantification of transcript numbers, *E. faecalis* OG1RF gDNA was used as template to generate standard curves. Relative quantification of gene transcripts was performed as previously described (67), using *gyrA* as the control gene.

### Analysis of proteinaceous metal chelators in wounds by ELISA

Naïve tissues were collected on the day of wounding by collecting the intact skin tissue removed by the biopsy punch. Tissues (naïve or from infected wounds) were excised and mechanically homogenized in 1ml metal-free PBS with 1U of protease inhibitor (Halt™ Protease Inhibitor, ThermoScientific).

The tissue was filtered using Flowmi 40 µm cell strainer tips (Sigma-Aldrich) to remove fibrous tissue. Samples were prepared following the instructions provided by each ELISA kit (all purchased from MyBioSource). For calprotectin (cat no. MBS7606640) and S100A7 (cat no. MBS763369) ELISAs, total protein content of the sample was determined using a BCA assay and samples were diluted to 1 mg ml^-1^ total protein content. The dilution factor for each sample was recorded for downstream analysis. For aTF ELISA (cat no. MBS729302), undiluted samples were used and for total transferrin ELISA (cat no. MBS2700560) a 1:10 (naïve skin tissue) or 1:100 (infected wound tissues) dilution of sample was used. For lactoferrin ELISA (cat no. MBS269515), samples were diluted 1:10 (naïve skin tissue) or 1:50 (infected wound tissues). ELISAs were performed according to manufacturer protocol with no deviation. Analyte available in undiluted samples were calculated based on dilution factors of sample and normalized to weight of tissue in grams.

### Statistical Analysis

All data sets were analyzed using GraphPad Prism 10 software. Data from multiple experiments conducted on non-consecutive days were collated. Data that falls within a Gaussian distribution were analyzed using either one-way or two-way ANOVA; however, a non-parametric test such as the Kruskal-Wallis or Mann Whitney was used if there is non-normality of data, both with appropriate comparison tests. Two-sided *p* values were calculated with *p* < 0.05 considered statistically significant.

## DATA AVAILABILITY

The authors confirm that the data supporting the findings of this study are available within the article and/or its supplementary materials.

## ACKNOWLEDGEMENT

This study was supported by R21 AI137446 awarded to J.A.L., American Heart Association (AHA) predoctoral fellowship (AWD 907592) awarded to D.N.B and AHA postdoctoral fellowship (AWD907586) awarded to L.N.L. D.N.B was also supported by T90 DE021990.

